# Phase separation of second prion domain of CPEB3: Insights from the aggregation and structural studies

**DOI:** 10.1101/2024.04.01.587532

**Authors:** Dhanya S Reselammal, Faina Pinhero, Arunima Sandeep, Vinesh Vijayan

## Abstract

The maintenance of long-term memory requires sustainable synaptic connections, mediated by the prion-like transformation of the translational regulator protein CPEB3 (Cytoplasmic Polyadenylation Element Binding protein isoform 3) in mammals. The N-terminal prion domain of CPEB3, composed of the two prion subdomains PRD1 and PRD2 has previously been demonstrated to perform a crucial role in imparting prion-like properties to the protein. We have already reported the amyloid-core of the first prion subdomain (PRD1) of the mouse CPEB3. Here, we have investigated the aggregation properties and the structural characteristics of the mouse PRD2 (mPRD2) in vitro. We found that the mPRD2 undergoes phase separation. Interestingly, the mPRD2 formed stable and amyloid-like solid condensates instead of the typical liquid condensate formation. Solid-state NMR and other biophysical studies revealed the existence of mixed secondary structures for mPRD2 in condensates. We propose that the distinct phase separation behaviour of the mPRD2 would be due to the conformational changes attributed to the pattern of the mPRD2 amino acid sequence, resulting in the formation of rigid and amyloid-like self-assembly.

## Introduction

Long-term memory persistence is achieved through synaptic plasticity, which requires synapse-specific local protein synthesis regulated at the translational level [1–3]. CPEB (Cytoplasmic Polyadenylation Element Binding protein) proteins are translational regulators which promote local protein synthesis in specific synapses by binding to the mRNAs that possess cytoplasmic polyadenylation elements in their untranslated regions [4]. In vertebrates and invertebrates, the CPEB proteins of different species are evolutionarily conserved to perform similar functions involving cell division, neuronal development, learning, and memory [5]. The prion-like transformations of the CPEB proteins such as ApCPEB (Aplysia), Orb2 (Drosophila), and CPEB3 (mammalian homolog) were found to be essential for the maintenance of long-term memory [6–11]. The disordered and aggregation-prone N-terminal domains of these proteins are responsible for their prion nature.

Structural studies on the *in vitro* amyloid fibrils of ApCPEB revealed that only the N-terminal prion domains form the fibril axis, and the C-terminal RNA binding domains project out of the fibril axis [12]. The rigid part of the ApCPEB fibrils has a mixed secondary structure composed of β-sheet, α-helical and random-coil conformations. Likewise, the Orb2 filaments extracted from Drosophila brain showed the presence of a hydrophilic amyloid core in the Q-rich region of its prion domain [13]. Furthermore, the Orb2 protein was recently demonstrated to undergo phase separation *in vitro* [14]. In the human isoform 2 of CPEB3 (hCPEB3), the specific segments formed by (1-300) and (200-450) regions of the prion domain were identified to promote the amyloid formation and liquid-liquid phase separation, respectively [15]. In the mouse homolog of CPEB3 (mCPEB3), the N-terminal prion-like domain has been compartmentalized into three subdomains, PRD1 (1-217), LMD (218-284), and PRD2 (285-449) (Fig. 1 *A*). Both PRD1 and PRD2 subdomains exhibit prion-like characteristics, while the LMD subdomain has been found to interact with the actin cytoskeleton [11]. We reported the *in vitro* amylogenic nature of the first prion domain (PRD1) of mCPEB3 [16]. The PRD1 fibrils consist of a β-rich core downstream to its Q-rich N-terminus. Sequence analysis has shown that the PRD1 and PRD2 regions of mCPEB3 share a remarkable 94% sequence similarity with the corresponding regions in hCPEB3 isoform 2 (Fig. S1). The sequence mismatches between the PRD1 regions in both organisms were quite random. Hence, it was not surprising that the individual PRD1 regions of both isoforms exhibited similar amyloid characters *in vitro* [15, 16].

**Figure 1.**
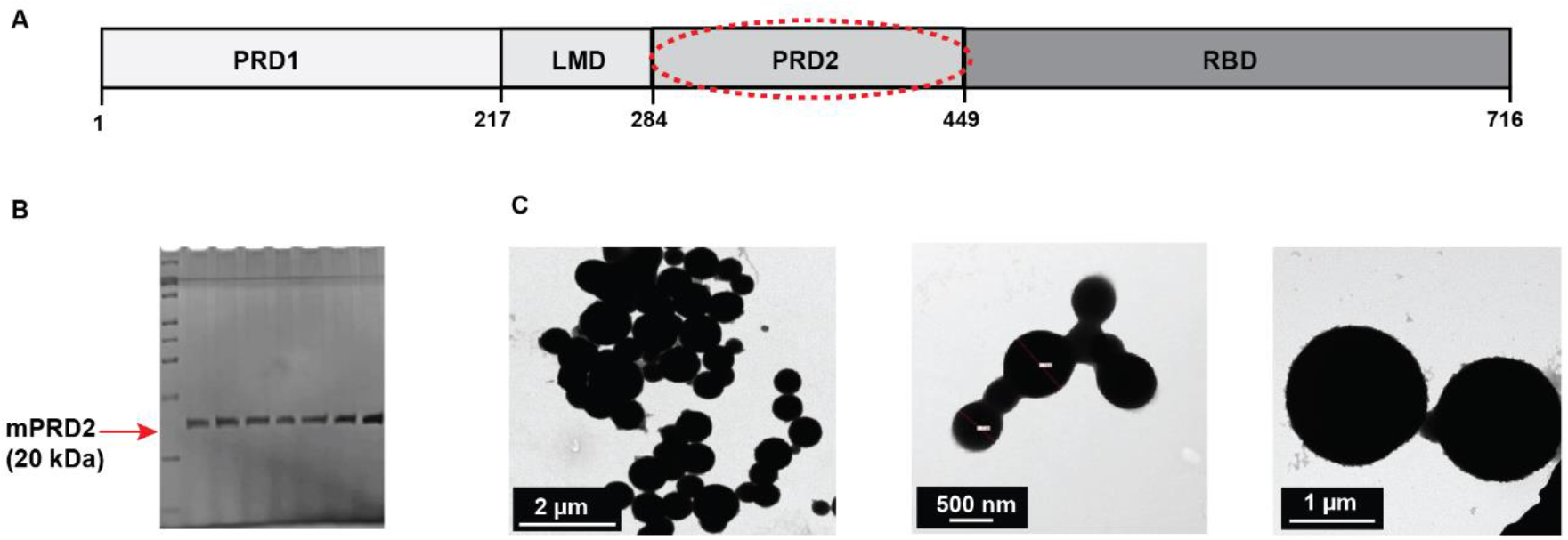
The recombinant mPRD2 refolds to form solid-like condensates. (A) Schematic diagram highlighting the domain organization of the full-length mCPEB3. The protein has a prion-forming N-terminus, which is subdivided into PRD1 (1-217), LMD (218-284), and mPRD2 (285-449) subdomains. The C-terminus contains the RNA binding domain (RBD, 450-716). The mPRD2 domain is highlighted with a dashed ellipsoid (red). (B) SDS-PAGE gel image shows an intense band corresponding to the pure recombinant mPRD2 protein at 20 kDa. The first lane shows the protein ladder. (C) TEM images of the mPRD2 solid-like condensates formed after refolding to the native buffer conditions.

The aggregation behavior and structural characteristics of the PRD2 domain of mCPEB3 (mPRD2) have not been explored thus far. In contrast, the PRD2 equivalent region in hCPEB3, spanning residues 284-417, has been demonstrated to be essential for the formation of liquid-like condensate [15]. We speculate that the structural role of the mPRD2 is critical to the mechanism of memory formation. Understanding the conformational differences of the PRD2 regions of the mammalian CPEB3 homologs would reveal the structural determinants contributing to the long-term memory establishment in mammals. Here, we investigated the role of mPRD2 in imparting the functional properties to mCPEB3. In contrast to hPRD2, which tends to form liquid-like condensates, our studies show that the mPRD2 preferably forms solid-like amylogenic condensates. Solid-state NMR studies on the condensates revealed the existence of mixed secondary structure conformations. Furthermore, we aimed to comprehend the structural changes in mPRD2 responsible for its distinct behavior in phase separation.

## Results

The recombinant mPRD2 domain of mCPEB3 undergoes condensate formation The mPRD2 domain (284-449) with a C-terminal histidine tag was expressed in *E- coli* cells (Materials and methods). Since the expressed mPRD2 was found to be associated with inclusion bodies, the protein was purified under denaturing conditions (8M urea, 50 mM tris-base, 100 mM NaCl, at pH 8). The purity of the protein was confirmed by SDS-PAGE, where a single protein band at 20 kDa in the gel image corresponded to the pure mPRD2 protein (Fig. 1B). Finally, the protein was gradually refolded under native conditions (50 mM Tris at pH 6.7, 1 mM DTT, 100 mM NaCl) through slow dialysis. During the dialysis process, the clear protein solution became turbid, indicating the formation of aggregates. Surprisingly, TEM images of the mPRD2 aggregates after refolding revealed the presence of protein condensates with a diameter in the range of 0.5-1 µm (Fig. 1C).

The morphology of the mPRD2 condensates closely resembled the recently reported solid-like condensates of the CPEB3 orthologs, Orb2A and Orb2B. [14] Condensates formed by truncated Orb2A and Orb2B proteins (Orb2AΔRBD and Orb2BΔRBD) transformed into mature fibrils over time. In contrast, the mPRD2 condensates remained stable, and no transition to a fibrillar state was observed. This behavior mirrored the characteristics of the full-length Orb2B, which also did not undergo further transformation into fibrils. Notably, the mPRD2 condensates were observed to merge upon contact with one another but did not fuse to form larger condensates.

Biophysical characterization of the mPRD2 condensates revealed its amylogenic nature The MALDI spectrum of the mPRD2 condensates was obtained by solubilizing in 50% acetonitrile. A high intense peak at 39.5 kDa, corresponding to the molecular weight of the mPRD2 dimer, and a less intense peak at 19.8 kDa, corresponding to the mPRD2 monomer was observed (Fig. 2 *A*). MALDI peaks indicated that the protein exists primarily as dimers in the solid-like condensates. The mPRD2 domain has only one cysteine residue, which is present at the C-terminal region of the protein. Therefore, the presence of dimers in the condensates suggests the formation of a disulfide bridge between the cysteine residues of the two protein monomers.

**Figure 2.**
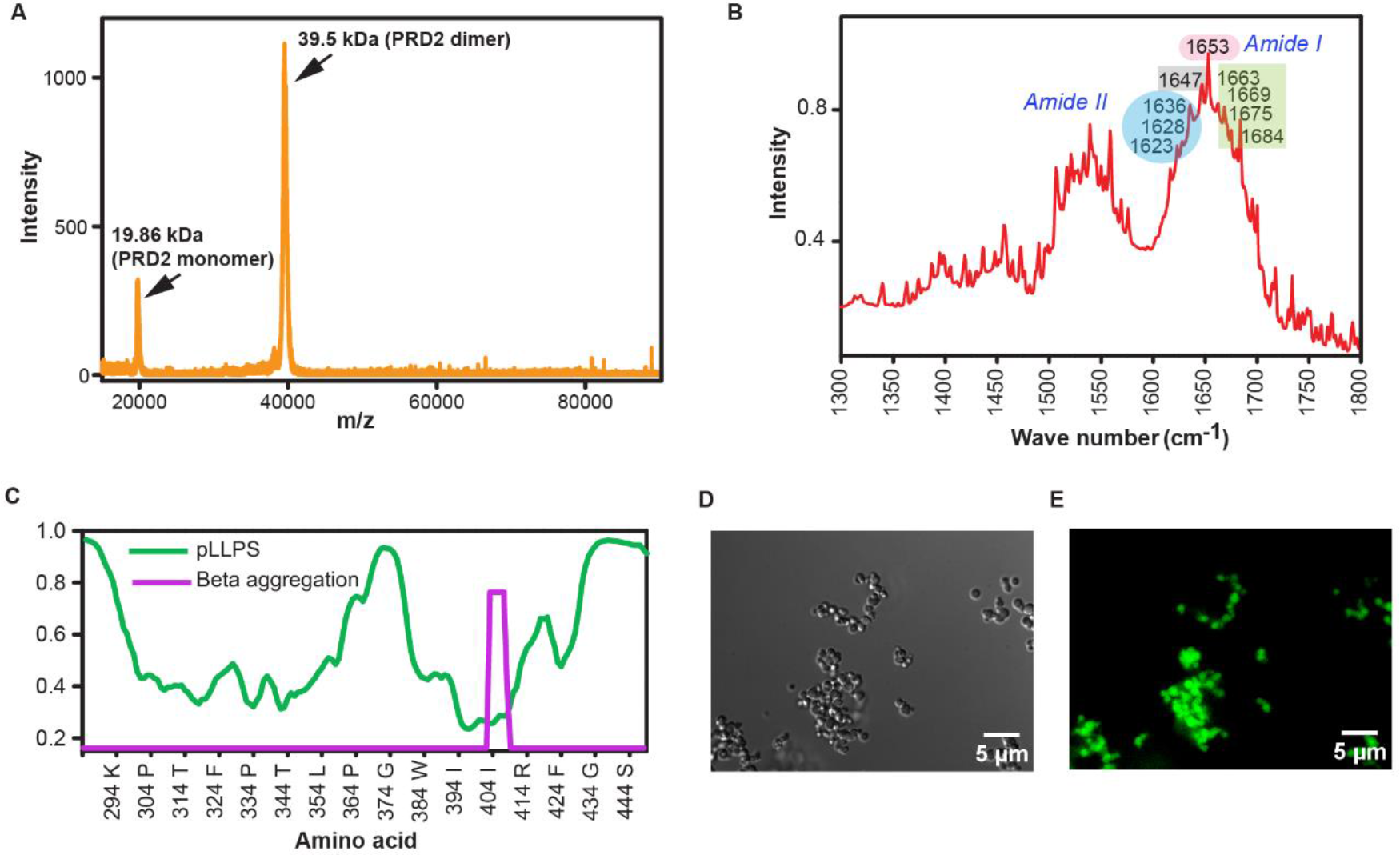
Preliminary characterization of the mPRD2 condensates. (A) MALDI spectrum of the mPRD2 condensates shows an intense peak corresponding to the mPRD2 dimer and a less intense peak corresponding to the monomer, suggesting that the mPRD2 oligomerizes largely as dimers in the condensates. (B) The FTIR (KBr pellet) spectrum of mPRD2 condensates washed and dried from D2O showed the presence of mixed secondary structures. Characteristic frequencies of the peaks in the amide I band corresponding to the β-strand (blue), random coil (grey), α-helix (magenta), and turn (green) conformations are indicated. (C) The plot (green trace) shows the residue-specific propensity of mPRD2 to undergo phase separation, as predicted using the FuzPred algorithm. The prediction showed probable phase separation propensity for specific stretches of residues (285-297, 361-379, 417-421, and 428-449). As predicted using the computer algorithm TANGO, the β-sheet aggregation propensity for the mPRD2 amino acid sequence is shown (violet trace). Non-zero β-aggregation propensity scores are observed for the residues in the stretch ^404^IMALNT^409^. (D) Representative DIC microscopic image of mPRD2 solid condensates. (E) Fluorescence microscopic image of the PRD2 condensates stained with Thioflavin T, on excitation at 440 nm shows bright fluorescence suggesting its β–rich characteristics.

Previous in-silico secondary structure analysis had predicted the presence of multiple helical motifs in mPRD2 (284–449) [11]. We performed a preliminary secondary structure characterization of the mPRD2 condensates using IR spectroscopy. To suppress the IR band from the solvent water (H2O) that overlaps with the amide-I IR band of the peptide bond, the condensates were repeatedly washed with D2O and dried [17]. The IR spectrum of the sample revealed mixed secondary structures within the condensates (Fig. 2 *B*). In the region of the broad amide I band, which is sensitive to the backbone conformation, there was an overlap of different secondary structures [18]. To identify these peaks, we utilized previously published mean frequencies for proteins in D2O [19]. The assigned peaks in the spectrum correspond to β-strand (in blue), random coil (in grey), α- helix (in magenta), and turn (in green) conformations (Fig. 2 *B*).

On analysing the mPRD2 amino acid sequence using the computer algorithm FuzPred, high phase separation propensities for certain regions: 285-297, 361-379, 417- 421, and 428-449 (Fig. 2 *C*, green trace) were observed [20]. But unlike the mostly observed liquid-like condensates formed by liquid-liquid phase separation, the mPRD2 condensates appeared to be solid-like. (DIC image in Fig. 2 *D*). The amyloid nature of the mPRD2 condensates was examined using the Thioflavin T (ThT) dye that specifically stains amyloids. Interestingly, the fluorescence microscopic image of ThT-stained mPRD2 condensates on excitation at 440 nm showed bright fluorescence upon ThT binding (Fig. 2 *E*), suggesting the presence of β-sheet rich amyloid regions within the condensates. This observation was reminiscent of the cellular membrane-less compartments formed by *Xenopus* balbiani bodies, which were reported to form amyloidogenic solid assemblies [21].

### TANGO predicted β aggregation-prone peptide motif shows high self-aggregation propensity

The ThT binding ability of mPRD2 condensates and the presence of β-sheet content in the FTIR spectrum clearly indicate amylogenic character. The prediction of the β-sheet-induced aggregation propensity of the mPRD2 sequence using the computer algorithm TANGO showed that a six-residue peptide motif (^404^IMALNT^409^) in the protein has non-zero β-aggregation propensity scores (Fig. 2 *C*, violet trace) [22]. Therefore, we examined the self-aggregation ability of the region by synthesizing an eight-amino acid long peptide (^403^NIMALNTR^410^), which contains the predicted motif. The peptide was synthesized using solid-phase peptide synthesis and was purified using HPLC (Fig. S2 *A*). As expected, the peptide was highly aggregation-prone. The peptide was found to be soluble only at pH 3.0. The residue-specific secondary structure propensity of the peptide was determined using solution-state NMR. To assign the peaks from the side chain and amide N-H regions of the peptide, we acquired ^1^H-^1^H TOCSY and ^1^H-^13^C HSQC spectra (Fig. S2 *B* and S2 *C-D*). Seven of the eight residues in the peptide sequence could be assigned. The Cβ of T409 could not be detected because of the overlap with the strong acetyl peak from the buffer. From the Cα and Cβ chemical shifts, the secondary chemical shift 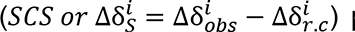 propensities of the peptide were calculated (the plot is shown in Fig. 3 *A*). Negative SCS values were obtained for six of the residues, suggesting the β-sheet propensity of the peptide. While the M405 exhibited a positive SCS value, which is a probable indication of a kink. Additionally, the peptide aggregation (200 µM) in 50 mM phosphate buffer (at pH 7) was monitored by incubating at 37 °C. We observed rapid fibril formation in 14 hours (TEM images, Fig. 3 *B*). However, the fibrils were found to convert further to longer-matured fibrils in 24 hours (TEM images, Fig. 3 *C*). Thus, the aggregation and structural studies confirm the amylogenic nature of the N403- R410 segment in mPRD2.

**Figure 3.**
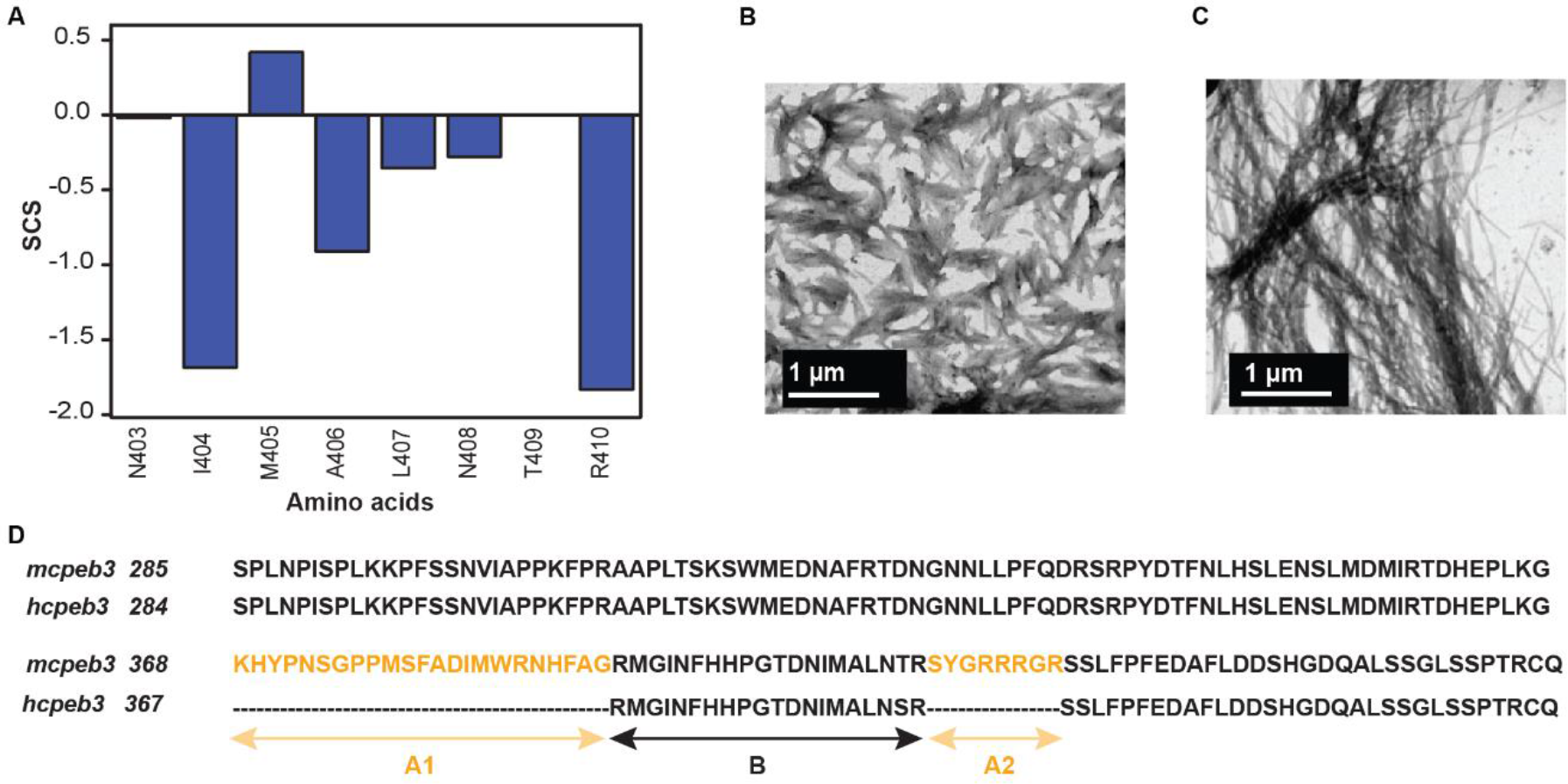
TANGO predicted β aggregation-prone peptide motif shows high self-aggregation propensity. Sequence alignment between mPRD2 and hPRD2 shows specific mismatches. (A) The SCS plot for the peptide fragment from mPRD2. Negative SCS values were observed for six of the residues, which suggests their β–sheet propensity. M405 showed a positive SCS value, which is the probable indication of a kink. SCS value for T409 was not obtained as it could not be assigned in the ^13^C-^1^H spectrum. (B) TEM image shows the fibrils of the peptide fragment from mPRD2 at a concentration of 200 µM in phosphate buffer at pH 7, formed within 14 hours of incubation at 37 °C. (C) The TEM image of the same peptide sample after 24 hours showed the presence of longer matured fibrils. (D) Sequence alignment between the mPRD2 equivalent domains of the mouse and human CPEB3 forms (mPRD2 and hPRD2). It shows two additional segments (A1 and A2) in the mPRD2 sequence (highlighted in orange) with a common middle segment (B).

### The difference in the phase separation behaviors of the mouse and human CPEB3 homologs; a sequence analysis

The mammalian CPEB3 homologs, mCPEB3, and hCPEB3 share ∼89% sequence identity. [15]. We analyzed the sequence alignment between the isoforms of mCPEB3 and hCPEB3, which had been previously studied for their aggregation and structural properties (Fig. S1) [11, 15, 16, 23]. The sequence alignment showed random mismatches in their PRD1 equivalent regions. In contrast, the sequence mismatches in their PRD2 equivalent regions; S285-Q449 (mPRD2) of mCPEB3, and S284-Q417 (hPRD2) of hCPEB3 are more specific (Fig. 3 *D*). The mPRD2 has two additional stretches of residues; (i) K368- G390 (in orange, denoted as A1) and (ii) S411-R418 (in orange, denoted as A2), connected by a common middle segment: R391-R410 (denoted as B). The two segments, A1 and A2 are missing in the hPRD2; hence this hCPEB3 isoform 2 is short by thirty-one residues. Whereas similar to the mPRD2 sequence, there are certain other human isoforms, which possess the A1 and A2 segments. This observation indicates the possibility of alternate splicing, which could have led to the deletion of the two segments in the genetic code of this specific human isoform 2.

Biophysical studies on the different segments of the prion domain of hCPEB3 showed that the region spanning the residues 254-426, corresponding to the hPRD2 region (S284-Q417), could undergo liquid-liquid phase separation. But the liquid-like condensates formed did not show any amylogenic character. In contrast, our findings have revealed the ability of the mPRD2 to form amyloid-like solid condensates. Hence, it is noteworthy to understand the structural characteristics of the mPRD2 solid condensates, which are responsible for their distinct phase separation behaviors.

### Monitoring the refolding of the mPRD2 domain from urea

To characterize the structural and dynamic properties of mPRD2 phase separation, we acquired a series of ^1^H-^15^N HSQC spectra at various urea concentrations, ranging from 8 M to 0 M of urea (Materials and methods). The ^1^H-^15^N HSQC spectrum of the mPRD2 protein under denaturing conditions (8 M urea, 1 mM DTT, 50 mM NaCl and 50 mM Tris at pH 6.7) exhibited relatively narrow amide resonances within a small spectral window of <1 ppm (within 7.8-8.8 ppm) (Fig. S3 *A*), confirming the dynamic and unstructured nature of the protein. Further, the denaturant was gradually removed by slow dialysis, and the HSQC spectrum was acquired at different urea concentrations. Interestingly, the mPRD2 sample showed phase separation at lower urea concentration.

To probe the refolding of the mPRD2 domain, we examined the chemical shift changes of the amide resonances from the protein with decrease in urea concentration. The ^15^N chemical shift difference plot (Fig. 4 *A,* cyan) of mPRD2 between 8 M urea and 3 M urea revealed an up-field shift for most of the residues. The segments W318-D321; L332-R338; M379-D381; W384-A389; R391-F396; N405-T409; S411-G416; L421-L429 showed an above-average chemical shift difference (Fig. 4 *A,* cyan). Noticeably, the W384-A389 in the A1 and N405-T409 in the B region exhibited relatively higher chemical shift differences compared to the rest of the segments. The prominent chemical shift changes in the A1 and B segments imply significant interactions in these regions during phase separation of the mPRD2. The observed peak shifts are nearly linear with the decrease in urea concentration (Fig. S3 *B-E*). Furthermore, with the decrease in urea concentration, a reduction in the number of peaks in the HSQC spectrum was also observed, possibly indicating the refolding of mPRD2 protein to produce phase-separated solid-like condensates. Eventually, the HSQC spectrum of mPRD2 in the buffer with nearly 0 M urea showed only a few peaks compared to the initial spectra (Fig. S3 *F*). This can be attributed to the remaining dilute phase concentration of mPRD2, which was found to be only 7 % of the initial concentration before phase separation.

**Figure 4.**
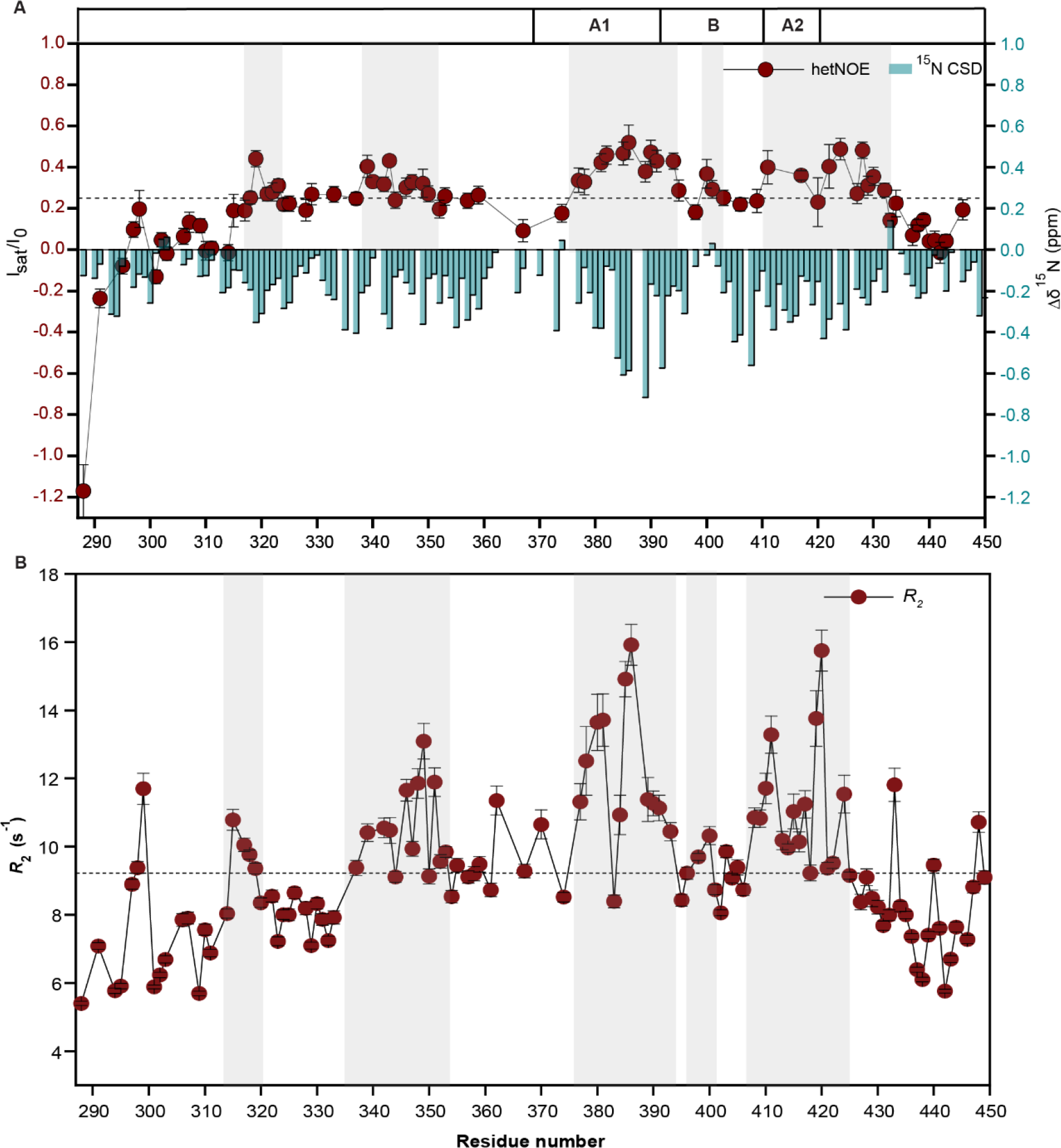
The solution NMR study of mPRD2 refolding from urea. (A) The overlaid plot of ^15^N chemical shift difference (^15^N CSD, cyan) between 3 M and 8 M urea containing mPRD2 and Het-NOE (dark red) of mPRD2 in 3 M urea at pH 6.7. The mean Het-NOE value is shown by dashed black line and the residues exhibiting above average Het-NOE values are highlighted in grey. The highlighted regions exhibit short segments that are involved in interactions mediating the phase separation of the mPRD2. (B) ^15^N transverse spin-spin (*R*2) relaxation rates of mPRD2 in 3 M urea at pH 6.7. The amino acid stretch with above average *R*2 value is highlighted in grey.

To study the backbone structural dynamics of mPRD2 in the presence of urea, we measured the ^1^H-^15^N Heteronuclear NOE (Het-NOE) and ^15^N transverse spin-spin (*R*2) relaxation rates of mPRD2 in 3 M urea (Fig. 4 *A,* dark red and 4 *B*). Generally, the cross peaks of amino acids with the intensity ratio (Isat/I0) below 0.6 suggests a rapid local conformational change of amino acids as well as the presence of protein backbone dynamics in the ps-ns timescale. [24] The average Het-NOE value of mPRD2 in 3 M urea was found to be 0.25, which further confirms that mPRD2 in 3 M urea is dynamically disordered. The residues at the N-terminus showed negative values for the NOE, as reported for most proteins. However, the C terminus of the protein was comparatively more rigid. Even though the protein is overall flexible, certain stretches of residues showed above average Het-NOE values (W318-A323; S339-L350; S378-N395; G400-N403; T409-S432), suggesting that these regions are more rigid compared to the rest of the protein sequence.

^15^N *R*2 of mPRD2 was measured in 3 M urea. ^15^N *R*2 values of residues spans from 4.68 s^-1^-15.92 s^-1^ with an average of 9.35 s^-1^. The *R*2 plot suggests a small degree of motional restrictions in the region involving residues S315-M319, D337-S353, M377-G393 and N408-F424. Interestingly, the pattern of clustering was observed to be similar to that obtained from the Het-NOE data. These short segments with elevated *R*2 values could be potential interaction sites that mediate phase separation. It is also to be noted that the stretch of amino acids in the A1 segment (M377-G393) showed maxima in both Het-NOE and *R*2 plot. It is probable that these segments make transient contacts during the phase separation process, which imparts partial rigidity.

### Structural analysis of the mPRD2 condensates using Solid-state NMR

As demonstrated using the solution NMR experiments, removing urea from mPRD2 suggested protein folding, leading to solid-like condensate formation. To further study the condensed phase of mPRD2, we employed solid-state NMR to examine the structural characteristics. The one-dimensional ^13^C CP spectrum showed intense signals, characteristic of different amino acids of the mPRD2 sequence (Fig. 5 *A*, red). Whereas the INEPT spectrum did not show any signals (Fig. 5 *A*, brown), suggesting that the condensates have high rigidity and hardly any flexible region. The two-dimensional ^13^C- ^13^C spin diffusion (SD) spectra of the condensates displayed resolved and sharp signals with narrow ^13^C line widths of ∼ 1 ppm, suggesting rigid and highly ordered structure (Fig. 5 *B*). Consistent with the IR data, solid-state NMR also revealed mixed secondary structures in the condensates. Amino acids with signature chemical shifts corresponding to β-sheet (labeled in blue), α-helix (magenta), and random coil conformations (green) were observed. Intra and inter-residue amino acid correlations were obtained from the SD spectra measured at two different mixing times (60 ms and 150 ms). In the overlaid SD spectrum (Fig. S4), sequential contacts for different stretches of residues in random coil conformation are denoted in green (P289-I290, R338-S339, etc.), those in α-helix in magenta (P375-G374, L429-D430, etc.) and those in β-strand in blue (G413-R414, N403-D402, etc.). The stretch of residues with their secondary conformation is represented in the mPRD2 sequence (schematic diagram, Fig. 6 *A*).

**Figure 5.**
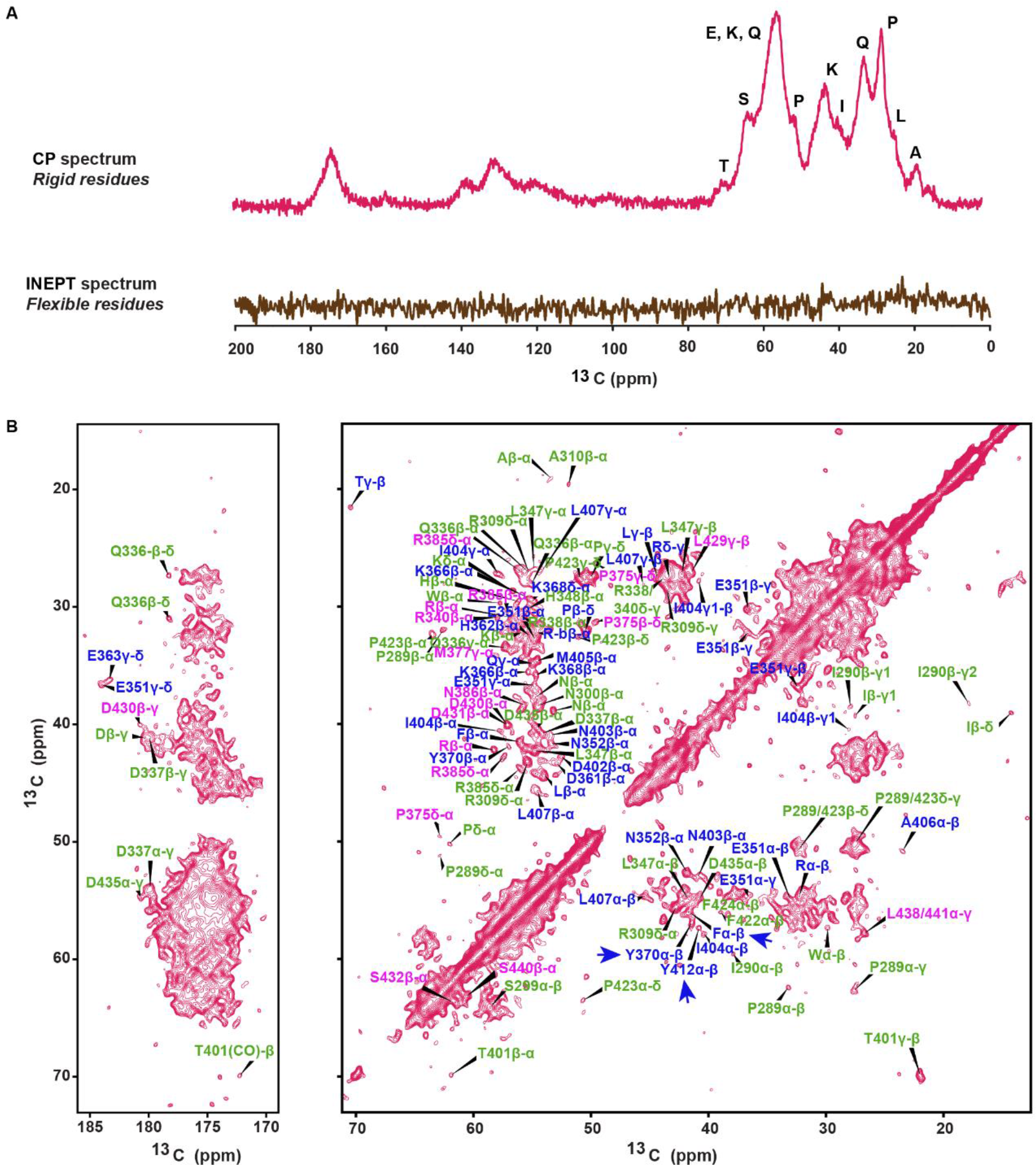
Solid-state NMR studies reveal the high rigidity of the mPRD2 condensates with mixed secondary structure. (A) One dimensional CP (red) and INEPT (brown) solid-state NMR spectra of the mPRD2 condensates acquired at 13 kHz MAS, and 5 °C in the 700 MHz NMR spectrometer. The ^13^C CP shows characteristic signals from various amino acids, whereas the ^13^C INEPT spectrum did not provide any signals. This suggested the high rigidity and lack of flexibility in the condensates. (B) Two dimensional ^13^C-^13^C SD spectrum (tmix = 60 ms) of mPRD2 condensates recorded at 18 kHz MAS, at -10 °C in 700 MHz spectrometer. The peak assignments for the residues in α-helix, β-sheet, and random coil conformations are labeled in magenta, blue and yellowish green respectively. The spectral windows correspond to the carbonyls (left) and the aliphatic carbons (right) of the protein. Only residue-type assignments with their characteristic conformations were denoted for the residues having an ambiguous sequential assignment. The assignments for the resonances from aromatic residues in β-sheet conformation are highlighted using blue arrows.

**Figure 6.**
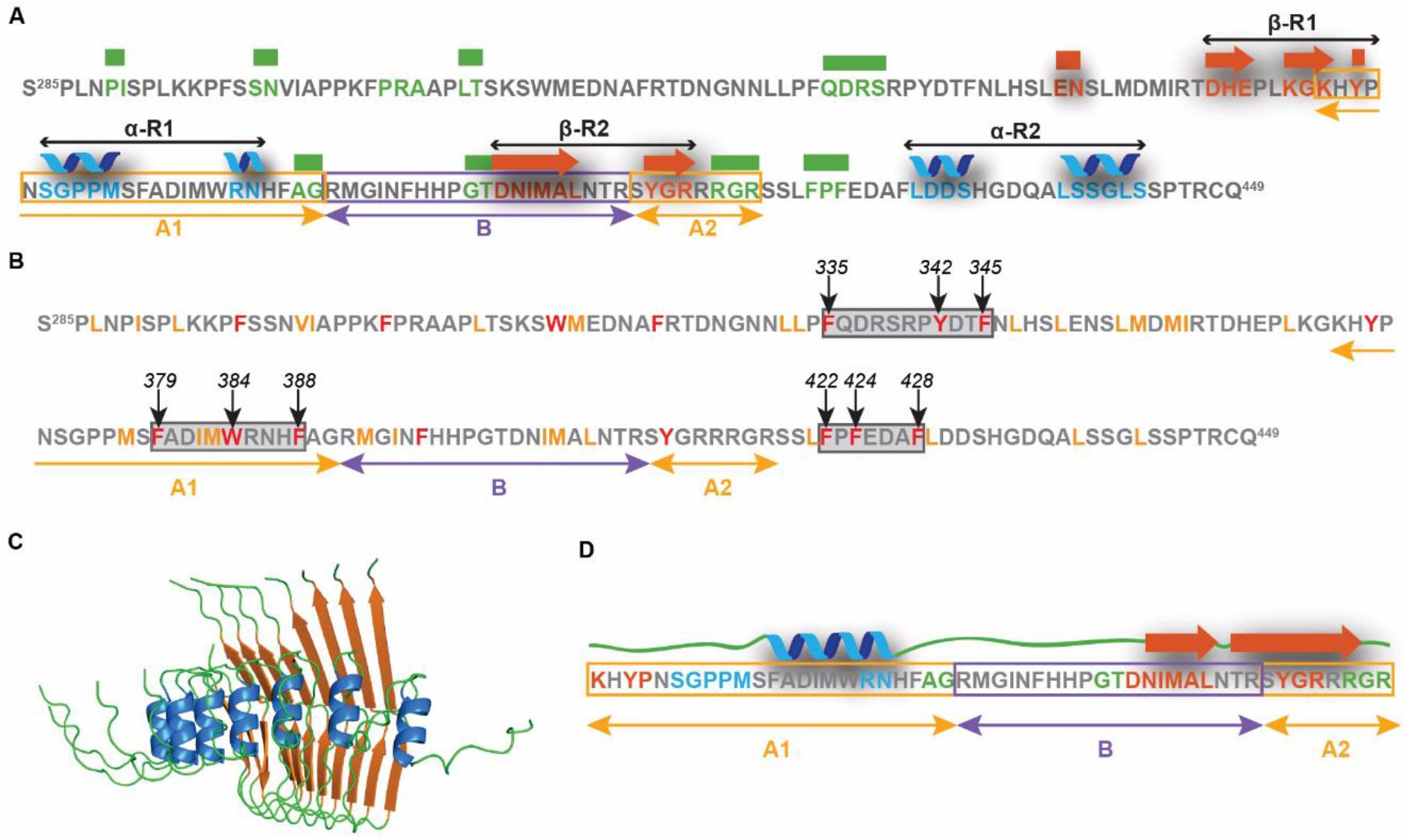
Structural analysis indicates the presence of segments with alternating random coil, β-strand, and α-helix conformations. (A) Schematic diagram highlighting the amino acid stretches in the mPRD2 sequence (165 amino acid long), which were identified to form α-helices (magenta), β-strands (blue), and random coil (green) conformations respectively. The additional A1 and A2 segments of mPRD2 are marked using orange rectangular boxes, and the common B segment is shown in a violet box. Solid-state NMR studies detected the presence of alternate β-sheet rich and α-helix-rich regions; β-R1, α- R1, β-R2, and α-R2. (B) The pattern of distribution of the aromatic and hydrophobic residues in the mPRD2 sequence. The aromatic and hydrophobic residues are highlighted in red and orange respectively. The closely spaced aromatic residues (F335-Y342-F345, F379-W384-F388, and F422-F424-F428) at three different regions are highlighted using grey-shaded boxes. (C) The AlphaFold2 predicted structure of the octamer of the 81 AA long (K368-R418) region of mPRD2, containing the two additional segments A1 and A2 (orange boxes) and the common B (violet box) segment of the mPRD2, as observed from the sequence alignment with the hPRD2. (D) The K368-R418 amino acid sequence, with the solid-state NMR, detected residues forming β-strand (blue), α-helix (magenta), and random coil (green) conformations are highlighted. A schematic showing the conformation of the corresponding residues as predicted by AlphaFold2 is shown on the top. It is observed that the experiment and the prediction do not overlap exactly. However, the solid- state NMR and the AlphaFold2 prediction indicated the presence of α-helical residues in the A1 segment and β-sheet residues in the B and A2 segments.

The partial resonance assignment of the SD spectrum provided evidence for the existence of alternating random coils, β-strands, and α-helices in the mPRD2 condensates (the schematic is shown in Fig. 6 *A*). Seven residues (S373-M377, R385-N386, in magenta) showed α-helix conformation in the A1 segment (in orange box, Fig. 6 *A*) of the mPRD2 sequence. Five residues (D402-L407, in blue) from the B segment (highlighted in a violet box) were detected to have β-sheet conformation. They also constitute the TANGO-predicted β aggregation-prone residues (I404-L407). Within the A2 segment of mPRD2 (in orange box, Fig. 6 *A*), we detected three residues (Y412-R414) adopting a β- sheet conformation. Thus, solid-state NMR studies unveiled the presence of alternating regions rich in β-sheet and α-helix; β-R1 (upstream to A1 segment), α-R1 (within A1 segment), β-R2 (within B segment), and α-R2 (towards PRD2 C-terminus). Most of the residues in these β-sheet rich and α-helix-rich regions were also previously found to have above-average Het-NOE values in the solution NMR measurement at 3 M urea, indicating more rigidity in these regions.

The clustering of aromatic residues and hydrophobicity of the residues between the aromatics have been demonstrated to be the sequence determinants for forming solid-like assemblies [25]. We observed the presence of resonances from specific aromatic residues in β-sheet conformation. Specifically, we unambiguously detected two tyrosines (Y370 and Y412). Additionally, there were phenylalanines with ambiguous sequential assignments (highlighted in blue arrows in the SD spectrum, Fig. 5 *B*). The mPRD2 sequence contains ∼10% of aromatic residues and ∼41% of hydrophobic residues; the pattern of their distribution in the sequence is shown in Fig. 6 *B*. The sequence also contains closely spaced aromatic residues (F335-Y342-F345, F379-W384-F388, and F422-F424-F428) at three different regions (highlighted using grey-shaded boxes).

However, a complete structural elucidation is required to obtain the three-dimensional structure of the mPRD2 condensates and to identify the molecular interactions contributing to the solid condensate formation.

Structural prediction using Alphafold2 showed mixed secondary structure for the octamer of mPRD2 fragment, A1-B-A2

Further, we predicted the three-dimensional structure of the mPRD2 oligomer using AlphaFold2, which predicts protein structures with the highest level of atomic accuracy [26]. For prediction, we used the amino acid stretch K368-R418, which constitutes the A1 and A2 segments and the common B segment of the mCPEB3. The octameric state of the K368-R418 stretch was predicted to have a β-sheet stacking structure (Fig. 6 *C*). AlphaFold2 also predicted the presence of mixed secondary structures in the A1-B-A2 segment; an α-helical motif (F379-H387) in the A1 segment, and two β- strands (N403-L407 and T409-R416) in the B and A2 segments (Fig. 6 *D*). Part of the A1 region was predicted to possess an α-helical segment (F379-H387); however, only a couple of residues were detected reliably by solid-state NMR (R385-N386) which were in α-helical and coincided with the prediction. Solid-state NMR also suggests α-helical secondary structure of A1 residues (S373-M377) that differ from AlphaFold2. AlphaFold2 has predicted additional regions in B and A2 to be in the β-strand. These residues were either unidentified or existed as a random coil when observed in solid-state NMR.

## Discussion

The complexity and uniqueness of memory lies in the molecular mechanisms of memory establishment. The mechanism of memory stabilization has not been explored completely. However, it is known that the prion state of CPEB3 plays a significant role in memory retention. The prion-states of the neuronal-specific isoforms of CPEB protein from divergent species are evolutionarily conserved to perform a common function in maintaining long-term memory [5, 27]. In contrast to the highly conserved RNA binding C-terminal domain, the Q/N rich N-terminal prion-like domain (PLD) varies across species. Unlike invertebrate orthologs, the mammalian CPEB3s have longer PLDs with a lesser number of glutamine residues [27]. The mouse and the human CPEB3 homologs share ∼89% sequence similarity [15]. Sequence alignment between the previously studied isoforms of mCPEB3 and hCPEB3 has revealed that the PRD1 equivalent regions show random mismatches [15, 16]. In contrast, the PRD2 equivalent regions exhibit specific mismatches, wherein two additional segments A1 (23 residues) and A2 (8 residues), are present in the mouse form (Fig. S1 and Fig. 4). We have already reported the amylogenic nature of the PRD1 of mCPEB3 [16]. Similarly, the PRD1 equivalent region of the hCPEB3 has also been demonstrated to produce amyloid fibrils with similar amyloid core forming region [28]. The PRD2 matching region in hCPEB3 (hPRD2) was interestingly found to undergo liquid-liquid phase separation [15]. The domain organization of CPEB3 with low complex regions and RNA-binding domains (RBDs) is reminiscent of certain RNA-binding proteins like fused in sarcoma (FUS) and TDP-43 that also undergo liquid-liquid phase separation [29]. In this study, we have investigated the aggregation and phase separation behavior of mPRD2 and compared our results with the already reported phase separation properties of hPRD2 [15].

The recombinant mPRD2 was soluble only under denaturing conditions. Upon refolding, the protein formed solid-like condensates, which were stable and amylogenic. Remarkably, the morphology of the mPRD2 condensates closely resemble that of the condensates formed by the CPEB3 orthologs Orb2A and Orb2B [14]. Solid-like phase-separated condensates are not unique and have been previously reported within cells as well. Examples include *Xenopus* balbiani bodies, centrosomes, nuclear pores, and yeast stress granules, all of which represent less dynamic biomolecular condensates within the cellular environment [21, 30, 31]. These biomolecular condensates differ from hydrogels and exhibit incomplete fluorescence recovery after photobleaching (FRAP), indicating their limited mobility [32].

While significant attention has been given to liquid-like condensates in which the components can be dynamically exchanged with the cellular environment, it is crucial to recognize that the dynamically arrested biomolecular condensates also form an essential class of cellular assemblies [25, 31–33]. The diversity in the protein assemblies ranging from viscous liquids to gels to solid-like functional amyloids, suggests that the material state can be finely tuned to perform specific physiological functions within cells [31]. The formation of solid-like condensates may serve as a mechanism by which the biomolecules achieve protection and tolerance from stress conditions such as heat, desiccation, etc [34, 35]. For example, Velo1 protein-induced amylogenic condensates play a role in forming balbiani bodies in dormant oocytes, providing a stable matrix for organelles [21]. The observation of the formation of amyloid-like solid mPRD2 condensates rather than liquid condensates formed by hPRD2 raises an important question as to the origin of the contradictory dynamics of the condensates.

Indeed, the protein sequence characteristics play a significant role in tuning the material state of the condensates [36, 37]. In addition, the sequence features might be evolutionarily selected to adopt specific functions within the cells [25, 38]. Analyses suggest that uniformly distributed aromatic and/or hydrophobic residues favor liquid-liquid phase separation and prevent aggregation [37, 39]. Specifically, the clustering pattern of aromatic residues and the amino acid composition between these aromatic residues are proposed as key determinants capable of fine-tuning the material properties of condensates [25]. The Velo1 PLD, which forms *in vitro* solid-like assembly, has been demonstrated to attain liquid-like condensate properties in the rationally designed PLD constructs that have either reduced aromatic clustering or on decreasing hydrophobicity between the aromatic residues [25]. In our case, higher Het-NOE and *R*2 values of residues at low urea concentration originate from peptide regions rich in aromatic residues. We also observed the presence of several aromatic residues in β-sheet conformation within the solid-state SD spectrum (highlighted in blue arrows, Fig. 5 *B*). The mPRD2 sequence contains several closely spaced aromatic residues, namely F335- Y342-F345, F379-W384-F388, and F422-F424-F428, situated in three different regions (highlighted using grey-shaded boxes, Fig. 6 *B*). Notably, the hPRD2, which forms liquid condensates, lacks the aromatic cluster F279-W384-F388 in the longest dissimilar A1 segment. Our solution-state NMR refolding studies suggested the involvement of the A1 segment of mPRD2 in the condensate formation. Though multiple features encoded in the mPRD2 sequence could contribute to the rigid and amyloidogenic character of the condensates, we expect that the A1 segment could probably be a key determinant in driving the solid-like phase separation of mPRD2.

Solid-state NMR studies on the mPRD2 condensates revealed its high rigidity, structural order, and lack of dynamic heterogeneity. In these condensates, we discovered the presence of mixed secondary structure characterized by alternating α-helical and β- strand forming regions (Fig. 6). Interestingly, previous research has demonstrated that disruption of α-helical assembly in TDP-43 inhibits liquid droplet formation [40]. Similarly, the β-stacking interactions in the mPRD2 condensates could disrupt the α-helical assembly and inhibit the formation of liquid condensate. This disruption may have contributed to developing rigid, amylogenic, and less dynamic mPRD2 condensates. Furthermore, prior studies have shown that SUMOylated mCPEB3 undergoes phase separation [41, 42]. Hence, it is likely that the SUMOylation of mCPEB3 exposes the interaction sites in the mPRD2 domain to facilitate condensate formation [43].

In conclusion, our studies prove that the mPRD2 region triggers the phase separation in mCPEB3. The specific dissimilarity in the amino acid sequences of the PRD2 equivalent regions of the mouse and human isoforms has resulted in their different phase separation behaviours. Complete structural characterizations of the hPRD2 liquid and mPRD2 solid condensates are required to reveal the critical interactions responsible for the conformational change. Nonetheless, the absence of the two additional segments, A1 and A2, in the hPRD2 sequence would be responsible for its liquid-liquid demixing, leading to increased protein dynamics for efficient control of biochemical reactions and probably enhanced mechanism of long-term memory retention.

## Materials and Methods

### Expression and purification of mPRD2 domain

The mPRD2 construct with a C-terminal His-tag was bought from Genscript Inc. in pET20b+ plasmid. The mPRD2 plasmid was transformed to E. coli Rosetta cells. The cells were cultured at 37 °C (225 rpm) in LB medium containing ampicillin resistance (1000 µg/ml) till the OD600 reached 0.6. The protein expression was induced with 0.8 mM IPTG for eight hours at 37 °C (220 rpm). Then the cells were harvested by centrifuging at 8000 rpm for 10 minutes and were stored at –20 °C for further use.

The cell pellets were dissolved in lysis buffer (50 mM Tris (pH 8), 1 mM PMSF, and 100 mM NaCl), and the cell lysates were sonicated for 15 minutes (30 seconds on, 30 seconds off, repeated 15 times) using a probe sonicator. It was then centrifuged at 8500 rpm, 4 °C for 50 min. Then, the supernatant was discarded, and the pellet was again dissolved in lysis buffer containing 1 % Triton. It was again centrifuged for 50 min at 8500 rpm and 4 °C. The pellet obtained was dissolved in Tris buffer containing 8 M urea. It was kept at 4 °C for 1 hour, slightly shaking, so that the pellet dissolved, and then centrifuged at 9000 rpm, 4 °C for 1 hr. The pellet was removed carefully, and the supernatant containing protein was collected. The supernatant with the his-tagged protein was loaded onto Ni-NTA superflow column pre equilibrated with equilibration buffer (8 M Urea, 50 mM Tris (pH 8), 50 mM NaCl, 0.03 % NaN3). It was incubated for column binding for 1 hour. Then the flow through was collected and the column was washed using 50 ml equilibration buffer. Then gradients of imidazole from 10 mM to 1 M in equilibration buffer were added to the column for eluting the protein.

### Expression and purification of ^15^N/^13^C labelled mPRD2 for solution-state and solid-state NMR studies

For the solution-state NMR structural studies, the mPRD2 was labelled with the NMR active isotopes ^15^N and ^13^C for conducting multidimensional experiments. The protein was expressed in ^15^N enriched minimal medium using the previously reported protocol for protein labeling [44–46]. The Rosetta cells containing the mPRD2 construct inserted in the pET20b+ plasmid were cultured in LB medium until the optical density (O.D600) was 0.6. The cells were pelleted and were resuspended either in the minimal medium containing ^15^NH4Cl (0.1% w/v) and D-glucose (0.4% w/v) to obtain the ^15^N uniformly single labelled protein or in a media containing ^15^NH4Cl (0.1% w/v) and ^13^C-labelled glucose (0.4% w/v) (Cambridge isotope Laboratories, Inc.) for ^15^N and ^13^C uniformly double labelled protein. The bacterial cells in the minimal media were grown at 37 °C until the O.D600 was increased by 0.1 units. Then the protein was overexpressed by inducing the cells with 1 mM IPTG and was incubated at 37 °C, shaking at 225 rpm for eight hours. The cells were pelleted by centrifuging at 9000 rpm and were stored at -20 °C. Subsequently, the cells were lysed and followed the same procedure of extraction and purification, which was carried out for the unlabeled mPRD2, mentioned above.

### Monitoring the refolding of PRD2 from urea

The mPRD2 protein at a concentration of 450 µM (3 ml) in the denaturing buffer (8 M urea, 50 mM Tris at pH 6.7, 1mM DTT and 50 mM NaCl) was slowly dialysed in a dialysis membrane to gradually reduce the urea concentration from the protein. With decreasing urea concentrations, aliquots of protein were taken to acquire ^15^N-^1^H HSQC NMR. All the solution NMR experiments were performed on Bruker Avance 700 MHz spectrometer equipped with a TXI probe at 298 K. The experiments were processed using NMRPipe software and analyzed using NMRFAM-Sparky. ^1^H-^15^N HSQC NMR measurements were acquired for the mPRD2 sample in the buffer containing urea concentrations of 7.5 M, 6.2 M, 5.7 M, 4.5 M, 3.7 M, 2.5 M, 2 M, 1.5 M, 0.9 M, 0.5 M and 0 M. The concentration of urea after each dialysis step was determined by calibrating the amide proton peak of urea at 5.78 ppm. The (^1^H-^15^N) heteronuclear NOE experiment was measured using the standard pulse sequence, hsqcnoef3gpsi. Interleaved experiments with and without proton saturation were conducted with a recycle delay of 4 s. The Het- NOE experiment was recorded with 256 and 2048 total points in the indirect ^15^N and direct ^1^H dimensions, respectively, with a sweep width of 18 and 14 ppm centered at 117.5 and 4.7 ppm, respectively. To measure the residue-specific ^15^N *R*2 relaxation rate constants, we used a ^1^H-^15^N HSQC-based 2D experiment with a relaxation compensated Car- Purcell-Meiboom-Gill (CPMG) scheme. A CPMG field of 550 Hz was used and each ^15^N *R*2 is made up of six interleaved relaxation delays. The variable relaxation delays were 3.4, 24.0, 51.4, 106.4, 171.6, and 209.4 ms. The experiment was recorded with 2048 complex data points in the acquisition and 1080 complex data points in the indirect dimension, respectively. The ^1^H-^15^N HSQC spectrum of 300 µM mPRD2 in 8 M urea was assigned using a series of triple resonance experiments such as HNCA, HNCACB and CBCA(CO)NH.

### TEM analysis of mPRD2 condensates

The mPRD2 condensates obtained by the removal of urea in the denaturing buffer (pH 6.7) through slow dialysis was centrifuged at 13000 rpm for 20 min and washed repeatedly. The pellets were adsorbed onto 400-mesh carbon-coated copper grids (Ted Pella) and stained with 0.4% uranyl acetate solution. The grids were analysed on a FEI TECNAI G2 Spirit BioTwin electron microscope at 120 kV.

### MALDI-TOF analysis of mPRD2 condensates

Equal volumes of mPRD2 condensates in water and TA50 (30:70 (v/v) of 0.1% Trifluoroacetic acid (TFA) in Acetonitrile to 0.1% TFA in water) were mixed to dissolve the condensates. Then, added with equal volumes of 2, 5-dihydroxybenzoic, acid (DHB) matrix (Sigma Aldrich). 1 µl was pipetted from each of the sample and was spotted on to a MTP 384 target plate ground steel (Bruker). MALDI-TOF analysis of the dried sample spots was carried out on a Bruker MALDI ultraflexTOF/TOF instrument, which provided the mass spectrum (relative intensity vs m/z) for the ions generated from the samples.

### Thioflavin T binding of mPRD2 condensates

The mPRD2 condensates were mixed with 0.5 mM Thioflavin T solution and the mixture was drop casted onto a glass slide. The sample drop was covered using a coverslip and was visualized using the fluorescence microscope Olympus IX83, with a 100x oil immersion objective. The images were acquired by exciting at 440 nm.

### FTIR spectroscopy of mPRD2 condensates

The IR spectrum of the mPRD2 condensates was obtained with a spectral resolution of 4 cm^-1^ on a Shimadzu IRPrestige-21 FTIR spectrometer by using the standard KBr pellet method.

### Solid phase peptide synthesis of the aggregation prone peptide from mPRD2

The eight amino acid long peptide from mPRD2 (^403^NIMALNTR^410^) was synthesised using automated PS3TM Peptide Synthesizer, utilizing[47]￼ Fmoc protected amino acids and the reagents for the synthesis were purchased from Sigma Aldrich. Dimethylformamide (DMF) was used as the solvent. 20% piperidine in DMF was used as the deprotector. 0.4 M N-methyl morpholine in DMF was used as the activator. The peptide was synthesized by attaching the C-terminal residue to the rink amide MBHA resin (Nova Biochem). HBTU ((2-(1H-benzotriazol-1-yl)-1, 1, 3, 3-tetramethyluronium hexafluorophosphate) (Sigma Aldrich) was used as the coupling agent. N-terminus of the peptide was kept unprotected.

After the synthesis, the resin was washed with dichloromethane (DCM) and dried. Finally, the side chains were deprotected and the synthesized peptide was cleaved from the resin under nitrogen atmosphere using the cleavage cocktail with the composition: 88% trifluoroacetic acid (TFA), 5% phenol, 5% water, and 2% triisopropyl silane (TIPS) for 150 minutes. The cleaved peptide was precipitated in ice cold ether, dried, and lyophilized from glacial acetic acid. Later the peptide was purified using reverse-phase high-performance liquid chromatography (HPLC) using C18 column in an HPLC instrument (Agilent Technologies). The pure peptide was lyophilized from water and was characterized using matrix-assisted laser desorption/ionization mass spectrometry.

### Solid-state NMR studies on mPRD2 condensates

The uniformly ^13^C, ^15^N labelled mPRD2 condensates were packed into a Bruker 3.2 mm thick-walled solid-state rotor. The solid-state NMR experiments were measured on Bruker 700 MHz NMR spectrometer equipped with 3.2 mm broad band MAS (Magic Angle Spinning) probe. One dimensional CP (cross polarization) and INEPT (Insensitive nuclei enhanced by polarization transfer) based spectra were acquired with 128 scans at 13 kHz MAS, 5 °C. The two dimensional [^13^C, ^13^C]-PDSD (proton driven spin diffusion) spectra at mixing times of 60 ms and 150 ms were obtained at 18 kHz MAS and at -10 °C with 260 scans, 1024 t1 points and 800 t2 points. The PDSD spectra were analyzed using Sparky software [48].

## Acknowledgments

D.S.R. and F.P. thank DST-INSPIRE and IISER Thiruvananthapuram for the PhD grants, respectively. V.V. thanks IISER Thiruvananthapuram and STARS/APR2019/BS/708/FS for funding.

## Author Contributions

D.S.R., and V.V. designed research; D.S.R., F.P., and A.S. performed research; D.S.R., F.P., A.S., and V.V. analyzed data; and D.S.R., F.P., and V.V. wrote the paper. ^1^D.S.R., and F.P. contributed equally to this work.

## Appendix A: Supplementary Material

Supplementary data to this article has been provided.

