## Supplementary material for "Phase separation of second prion domain of CPEB3: Insights from the aggregation and structural studies"

**SUPPLEMENTARY INFORMATION**

Supplementary Methods     …........................................... 2

Supplementary Figure 1      …........................................... 2

Supplementary Figure 2	…........................................... 12

Supplementary Figure 3	…........................................... 13

Supplementary Figure 4	…........................................... 15

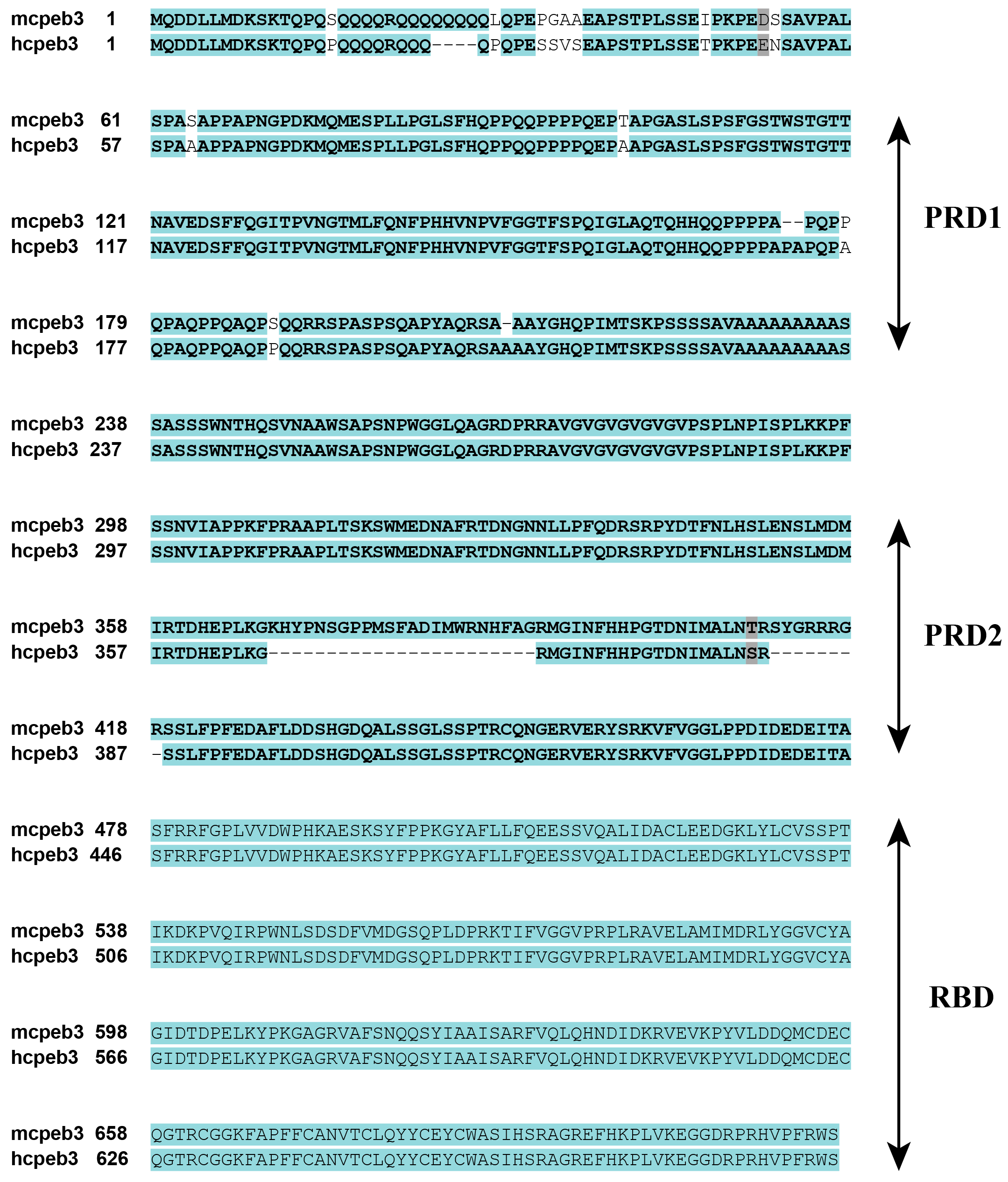

Supplementary Fig. S1. The sequence alignment between the mouse and human homologs of CPEB3 proteins, mCPEB3 (accession: AAQ20843) and hCPEB3 (accession: NP_001171608). The alignment showed 94 % similarity and 91 % identity for the PRD1 equivalent regions, 94 % similarity and 93 % identity for the PRD2 equivalent regions, and 100 % identity for the RNA binding C-terminus.

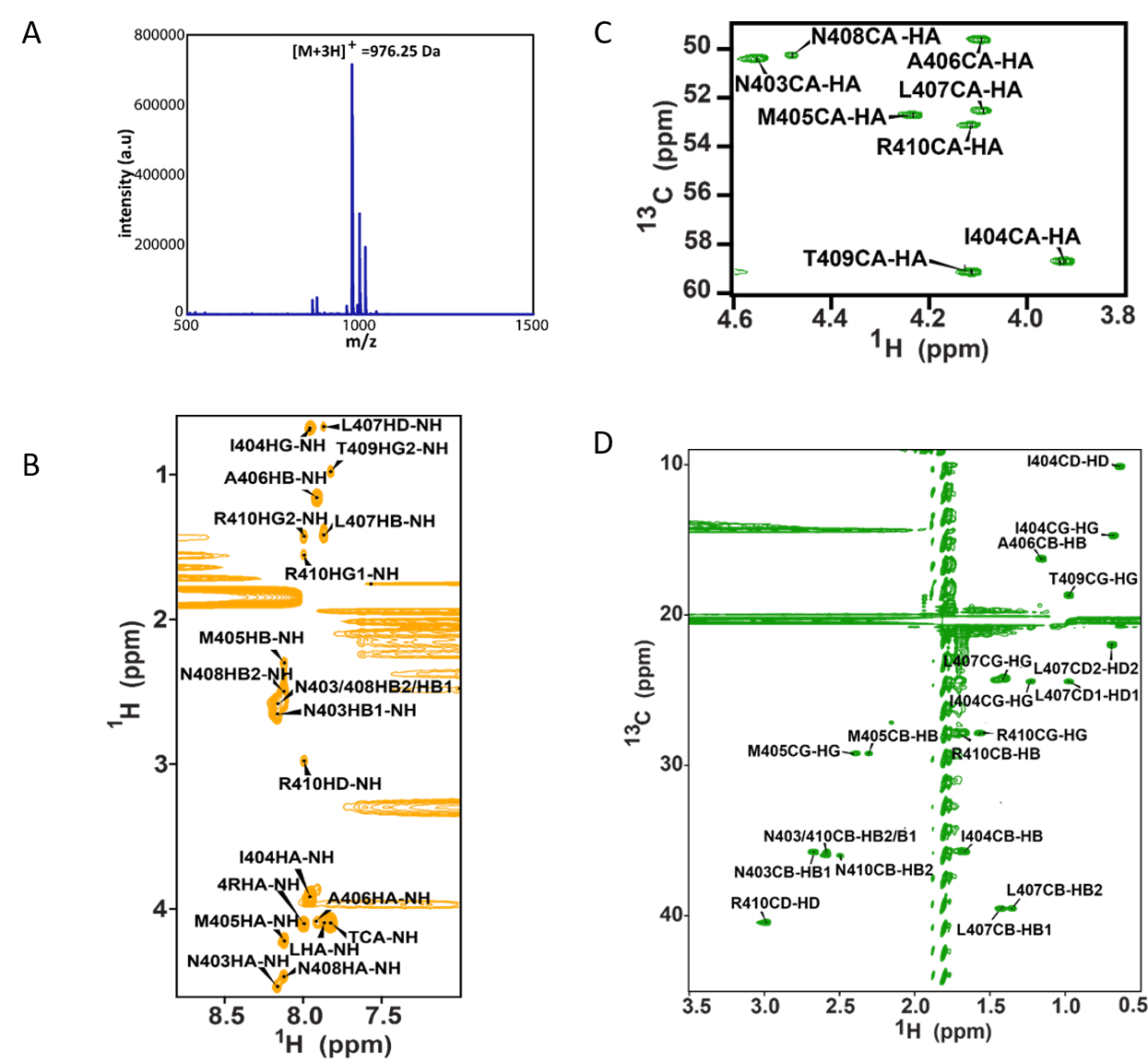

Supplementary Fig. S2. (A) The MALDI spectrum of the purified synthetic peptide from PRD2 (N403-R410) of 973 Da, shows intense peak corresponding to the [M+3H]^+^ ion. (B) The assigned ^1^H-^1^H TOCSY NMR spectrum of the peptide from PRD2 at a concentration of 1 mM measured with 16 scans and 512 indirect points. The peptide was found soluble only at pH 3, hence dissolved in 50 mM sodium acetate buffer at pH 3 with 10 mM NaCl and 0.03 % NaN3. ^1^H-^1^H TOCSY spectrum shows the cross peaks due to proton-proton through bond correlations from each of the amino acid in the peptide. (C-D) The assigned heteronuclear ^1^H-^13^C HSQC spectrum of the peptide from PRD2 (1 mM), measured with 32 scans and 256 indirect points. The spectral region in (C) shows the Cα-Hα correlations, and that in (D) shows the Cβ-Hβ, and Cγ-Hγ correlations. The NMR measurements were acquired in the 700 MHz Bruker NMR spectrometer equipped with cryoprobe.

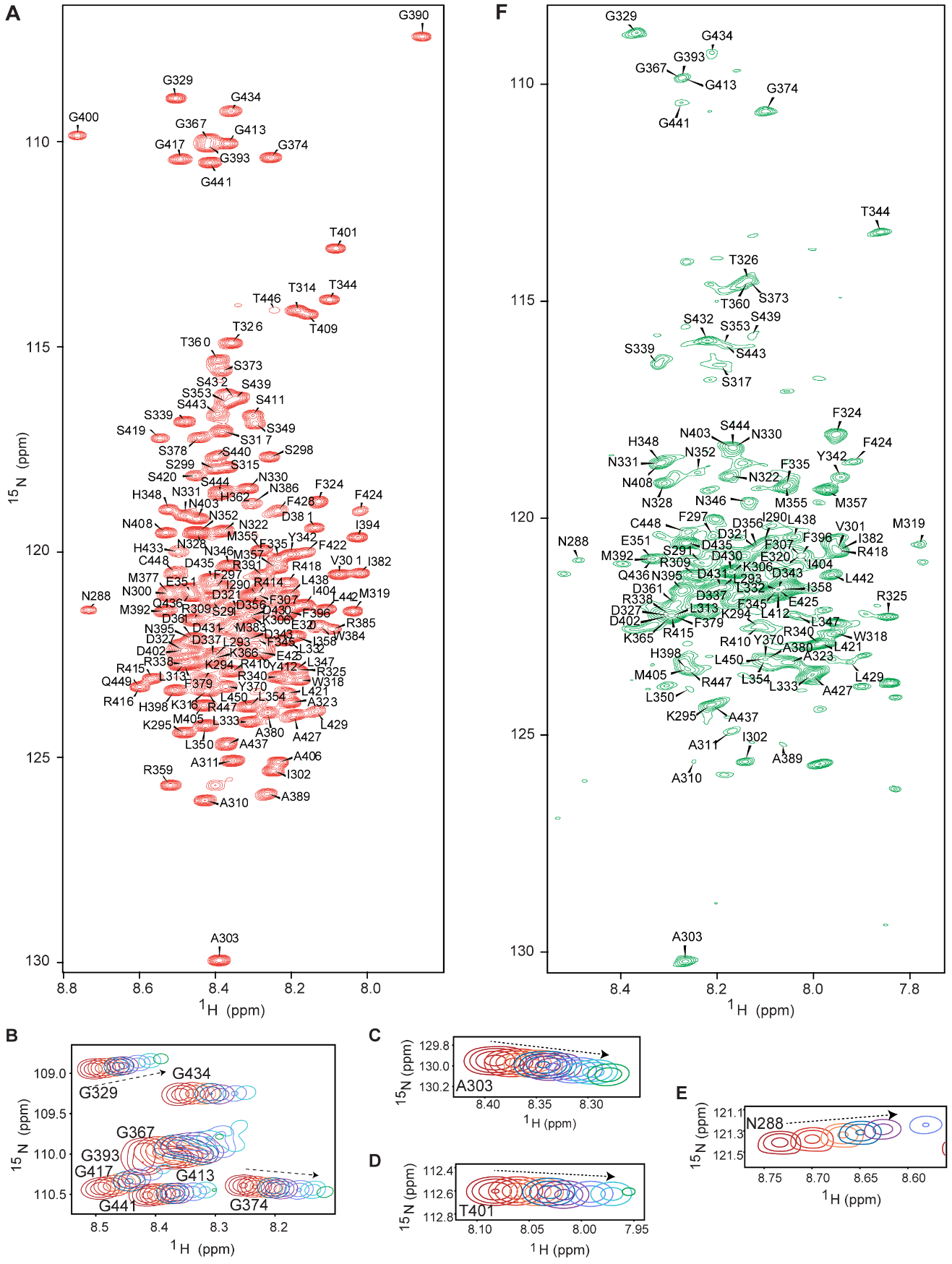

Supplementary Fig. S3. (A) The assigned ^1^H-^15^N HSQC spectrum of the mPRD2 protein under denaturing conditions (8 M urea, 1 mM DTT, 50 mM NaCl and 50 mM Tris at pH 6.7 and 0.03 % NaN3). It shows relatively narrow amide resonances within a small spectral window of <1 ppm (within 7.8 to 8.8 ppm). This confirms the unstructured and dynamic nature of the protein. For the resonance assignment of the peaks in ^1^H-^15^N HSQC spectrum of mPRD2, 3D triple resonance NMR experiments such as HNCA, HNCACB and CBCA(CO)NH (Materials and methods) were measured using uniformly ^15^N, ^13^C labelled mPRD2 sample containing 6 M urea, 50 mM NaCl, 0.03 % NaN3 and 50 mM Tris at pH 6.7. To monitor the conformational change during refolding, a series of HSQC spectra was acquired with decreasing concentrations of urea. The peaks in the ^1^H-^15^N HSQC spectra showed a gradual shift in their chemical shift positions during the refolding, and the observed peak shifts are effectively linear with the decrease in urea concentration; the spectral region of the Glycine residues (B), A303 (C), T401 (D) and N288 (E) are shown. (F) The ^1^H-^15^N HSQC spectrum of the mPRD2 protein obtained after slow dialysis to the buffer with nearly 0 M urea (1 mM DTT, 50 mM NaCl and 50 mM Tris at pH 6.7 and 0.03 % NaN_3_. It showed a fewer number of peaks with significant broadening compared to the initial spectrum (Supplementary Fig. S3A), indicating the formation of phase-separated solid condensates.

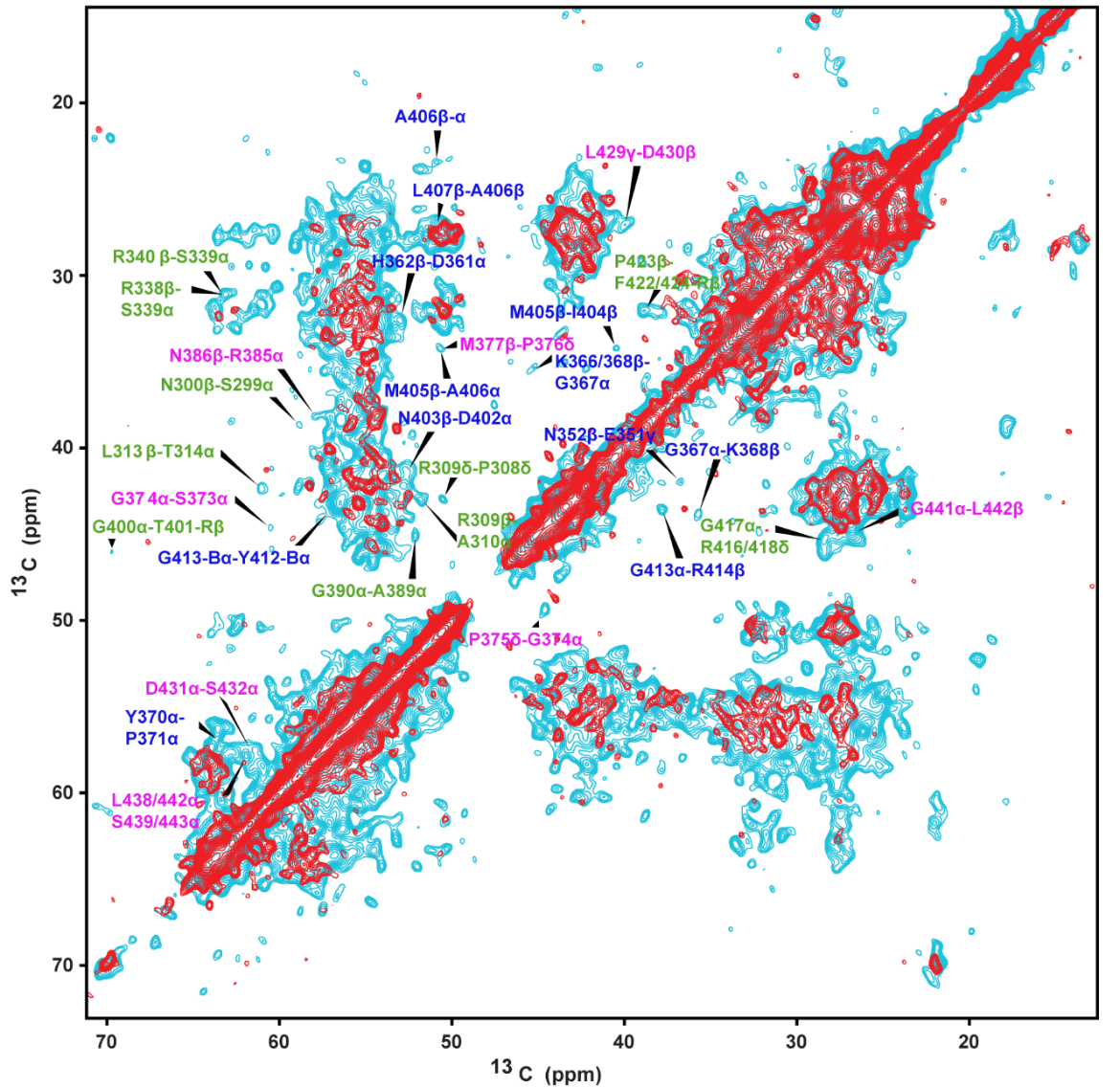

Supplementary Fig. S4. Overlaid ^13^C-^13^C SD spectrum of mPRD2 condensates recorded at the mixing times of 60 ms (red) and 150 ms (cyan) providing intra and inter residue correlations. Cross peaks due to inter-residue contacts of adjacent amino acids are marked in the spectrum. The secondary structure conformations of the correlated residues are highlighted in magenta, blue and green respectively for the α-helix, β-sheet and random coil. The measurements were conducted at 18 kHz MAS, at -10 ºC in 700 MHz spectrometer.
